# Using background noise to improve sound localization following simulated hearing loss

**DOI:** 10.1101/673806

**Authors:** Lindsey Ryan-Warden, Eva Ng, Peter Keating

**Author notes:** Correspondence to: Peter Keating, Ear Institute, University College London, 332 Grays Inn Road, Kings Cross, London WC1X 8EE, UK.

## Abstract

Many listening abilities become more difficult in noisy environments, particularly following hearing loss. Sound localization can be disrupted even if target sounds are clearly audible and distinct from background noise. Since subjects locate sounds by comparing the input to the two ears, sound localization is also considerably impaired by unilateral hearing loss. Currently, however, it is unclear whether the effects of unilateral hearing loss are worsened by background noise. To address this, we measured sound localization abilities in the presence or absence of broadband background noise. Adult human subjects of either sex were tested with normal hearing or with a simulated hearing loss in one ear (earplug). To isolate the role of binaural processing, we tested subjects with narrowband target sounds. Surprisingly, we found that continuous background noise improved narrowband sound localization following simulated unilateral hearing loss. By contrast, we found the opposite effect under normal hearing conditions, with background noise producing illusory shifts in sound localization. Previous attempts to model these shifts are inconsistent with behavioural and neurophysiological data. However, here we found that a simple hemispheric model of sound localization provides an explanation for our results, and provides key hypotheses for future neurophysiological studies. Overall, our results suggest that continuous background noise may be used to improve sound localization under the right circumstances. This has important implications for real-world hearing, both in normal-hearing subjects and the hearing-impaired.

**Significance Statement:** In noisy environments, many listening abilities become more difficult, even if target sounds are clearly audible. For example, background noise can produce illusory shifts in the perceived direction of target sounds. Because sound localization relies on the two ears working together, it is also distorted by a hearing loss in one ear. We might therefore expect background noise to worsen the effects of unilateral hearing loss. Surprisingly, we found the opposite, with background noise improving sound localization when we simulated a hearing loss in one ear. A simple hemispheric model of sound localization also helped explain the negative effects of background noise under normal hearing conditions. Overall, our results highlight the potential for using background noise to improve sound localization.

## Introduction

In everyday environments, a key challenge for the brain is to locate sounds of interest in the presence of background noise. Background noise can disrupt sound localization by reducing the audibility of target sounds (Good and Gilkey, 1996; Abouchacra et al., 1998; Lorenzi et al., 1999a, b; Brungart et al., 2005; Kopco et al., 2010; Kerber and Seeber, 2012; Lingner et al., 2012; Wood and Bizley, 2015). However, sound localization can be affected even if targets are clearly audible and distinguishable from concurrent sounds. In such circumstances, the perceived location of a target sound is typically pushed away from the location of a concurrent sound (Suzuki et al., 1993; Canevet and Meunier, 1996; Getzmann, 2002; Best et al., 2005; Lee et al., 2009; Reed and van de Par, 2015). Previous attempts to explain this have relied upon a neural map of space that contains an array of neurons (or ‘spatial channels’) tuned to different spatial locations (Suzuki et al., 1993; Best et al., 2005). However, these models predicted greater pushing effects between sounds that are closer together, which is inconsistent with previous behavioral data (Best et al., 2005). They are also inconsistent with neurophysiological studies in mammals, which have shown that neurons in each hemisphere are broadly tuned to a single side of space (Brugge et al., 1998; Furukawa and Middlebrooks, 2001; McAlpine et al., 2001; Grothe et al., 2010; Mokri et al., 2015).

Sound localization is additionally impaired by hearing loss, with the nature of impairment depending on the type of hearing loss experienced (Lorenzi et al., 1999a; Best et al., 2011; Dobreva et al., 2011; Akeroyd, 2014; Brungart et al., 2014). Under normal hearing conditions, sound localization relies on the relative timing and level of input to the two ears (Interaural Time Differences: ITDs; Interaural Level Differences: ILDs)(Middlebrooks and Green, 1991). Binaural sound localization is therefore particularly vulnerable following unilateral hearing loss, which alters and degrades these binaural spatial cues (Slattery and Middlebrooks, 1994; Hawley et al., 1999; Van Wanrooij and Van Opstal, 2004; Rothpletz et al., 2012; Agterberg et al., 2014; Firszt et al., 2017; Nelson et al., 2018). Key insights into this problem have been obtained by using earplugs to simulate a hearing loss in one ear (Slattery and Middlebrooks, 1994; Van Wanrooij and Van Opstal, 2007; Kumpik et al., 2010; Irving and Moore, 2011; Strelnikov et al., 2011; Keating and King, 2013; Keating et al., 2016; Asp et al., 2018). These studies have shown that sound localization is initially disrupted but improves as individuals adapt. Adaptation can be achieved by learning to locate sounds using the input provided to the unoccluded ear (Kumpik et al., 2010; Keating et al., 2013; Keating et al., 2016). This is possible because the head and ears filter sounds (i.e. change their spectrum) in a direction-dependent way (Carlile et al., 2005). However, when the energy of a sound is concentrated in a narrow range of frequencies, these monaural spectral cues are unavailable. In such circumstances, subjects can adapt by instead learning to reinterpret the altered binaural cues (Gold and Knudsen, 2000; Keating et al., 2015; Keating et al., 2016). Nevertheless, when adult humans wear an earplug for prolonged periods of everyday life, they find it difficult to do so (Kumpik et al., 2010).

Although background noise is a common feature of everyday life, previous studies of unilateral hearing loss have tested sound localization in quiet environments. For many auditory abilities, individuals with hearing loss typically experience greater difficulty in the presence of background noise (Moore, 1996; Bronkhorst, 2000; Lorenzi et al., 2006; Helfer and Freyman, 2008; Akeroyd, 2014). Consequently, we might expect individuals with simulated unilateral hearing loss to be particularly vulnerable to the effects of background noise. Surprisingly, however, we found the opposite. Following simulated unilateral hearing loss, background noise improved narrowband sound localization. A simple neurophysiological model also provided insight into how background noise might affect behavior.

## Materials and Methods

### Subjects

10 subjects (2 males and 8 females), aged 21-35, participated in the study. All participants provided informed consent and were reimbursed for their time. Participants underwent audiometry to confirm normal hearing prior to testing. Only one participant had prior experience of a sound localization task. All procedures were approved by the University College London Research Ethics Committee.

### Apparatus

Sound localization was tested in a double-walled anechoic chamber. Stimuli were presented from a semi-circular array of nine loudspeakers (Canton Plus XS.2; Computers Unlimited London), located in the front hemifield, with a radius of 1.2m. Participants were seated on a stool in the center of the speaker array, facing a central speaker at 0° azimuth. The height of the stool was adjusted to ensure the participant’s head was level with the speaker array; a chin rest was used to minimize head movements.

Stimuli were generated using the Psychophysics Toolbox (Brainard, 1997) for Matlab(The Mathworks, Natick, MA), sent to a MOTU 24io sound interface (MOTU, Cambridge, MA), amplified (MA1250; Knoll Systems, Point Roberts, WA) and presented via the appropriate loudspeaker. On each trial, a target sound was presented from a single loudspeaker. Participants indicated the perceived location of this target by clicking on a graphical user interface (GUI; generated in Matlab), which was presented on a screen located just below the central speaker. The GUI contained an illustration of the speaker array, which allowed participants to click on different response locations.

### Procedure

#### Familiarization

Before testing began, participants were given the opportunity to familiarize themselves with the sound localization task under normal hearing conditions. Throughout this familiarization process, the background of the graphical user interface changed color to indicate whether a behavioral response was correct (green) or incorrect (red). Following incorrect trials, the same target stimulus was presented again and participants were given a second opportunity to respond (referred to as ‘correction trials’). Following consecutive incorrect responses, the target stimulus was presented continuously for > 2s before participants were asked to respond (referred to as ‘easy trials’).

#### Testing

Each participant was tested in the presence or absence of background noise, with either normal hearing or a simulated hearing loss in one ear (see below), for a total of 4 unique test conditions (i.e. normal hearing in quiet; normal hearing in background noise; earplug in quiet; earplug in background noise). Within each session, participants were tested for 15 mins on each of these conditions (~400 trials), with the testing order randomized across participants. Participants completed two such sessions on different days, each of which lasted approximately 1hr, and were given short breaks between conditions. During these test sessions, participants were not given any feedback on their performance; correction trials and easy trials (see above) were also turned off.

#### Stimuli

Target sounds consisted of broadband noise (0.5-20kHz) or pure tones (1, 2, 4 or 8 kHz), which were identical to those used in previous work (Keating et al., 2016). Broadband and narrowband sounds were presented with equal probability. Broadband targets had either a flat or random spectrum, with random spectra produced by adding a random vector to the logarithmic representation of the source spectrum on each trial. This random vector was smoothed to remove spectral transitions > 3 cycles/octave and had an RMS of 10 dB (Keating et al., 2016). All target sounds were 100ms in duration (including 10ms cosine ramps), generated with a sampling frequency of 48 kHz, and presented at 56-77 dB SPL in increments of 7 dB. The intensity, type and location of target stimuli were randomly interleaved across trials. Background noise consisted of broadband noise (0.5-20kHz) with a flat spectrum presented continuously at the midline at 56 dB SPL. By setting the background noise to this level, we were able to vary target intensity (corresponding to different signal-to-noise ratios: SNRs) across a relatively wide range (21 dB), whilst ensuring that all targets were clearly audible (SNRs ≥ 0). Previous studies of unilateral hearing loss have shown that, if target intensity is not varied, subjects can locate sounds using the sound level in the better ear (Van Wanrooij and Van Opstal, 2004).

#### Simulated unilateral hearing loss

To simulate a hearing loss in one ear, participants wore a foam earplug (EAR classic) in one ear (Keating et al., 2016). Each participant wore the earplug in the same ear across sessions, but different participants wore the earplug in either the left (n=5) or right (n=5) ear. For each subject, conventional audiometry was performed in the absence and presence of the earplug. This allowed us to assess the attenuating effects of the earplug, which were very similar to those observed in previous studies (Kumpik et al., 2010; Keating et al., 2016).

### Statistical Analyses

To provide an overall measure of performance, we calculated the average magnitude of errors made by each participant across trials. These values were calculated separately for different hearing conditions (normal or earplug), background noise conditions (quiet or background noise), sound levels and stimulus types (broadband or narrowband). The statistical significance of these fixed effects was then assessed using mixed-effects ANOVAs, with subject as a random effect, followed by appropriate post-hoc tests corrected for multiple comparisons.

To facilitate comparison with previous work (Kumpik et al., 2010; Keating et al., 2016), and to help understand the reasons for subjects’ errors, we also performed linear regression to calculate subjects’ bias values as well as the slope of the relationship between stimulus and response. Perfect performance on our task produces a bias of 0 and a slope of 1, with deviations from these values reflecting errors in sound localization. Following previous work (Kumpik et al., 2010), we restricted this analysis to stimulus locations ≤30° from the midline. As above, these measures were calculated separately for different hearing conditions, background noise conditions, sound levels and stimulus types; significance was then assessed using mixed-effects ANOVAs and post-hoc tests corrected for multiple comparisons.

To test the predictions of the hemispheric model (see below), we calculated the mean response for each stimulus location in quiet and assessed how it changes in the presence of background noise. Since this requires separate measures for each stimulus location, data were pooled across sound level. For normal hearing conditions, we found that our data were symmetric around the midline (i.e. data for the left and right sides of space were opposite in sign but were otherwise very similar). To test this symmetry directly, our statistical model therefore included hemifield (left or right side of space), eccentricity (distance from the midline), and stimulus type (narrowband or broadband) as factors; our dependent variable was the extent to which sounds are pushed away from the midline. The significance of these fixed effects was then assessed using mixed-effects ANOVAs with subject as a random factor, followed by post-hoc tests corrected for multiple comparisons. Data for normal hearing and plugged conditions were analysed separately. All statistical analyses were conducted in Matlab.

### Hemispheric Model

We implemented a simple and popular model of sound localization (McAlpine et al., 2001; Keating et al., 2015) and applied it to previously published neurophysiological data (Furukawa and Middlebrooks, 2001). Full details of how these neurophysiological data were collected have been described previously by the authors. Briefly, neuronal responses were recorded in anaesthetized cat A2 (right hemisphere). Target sounds were presented at different locations (−160 to 160 in increments of 40) either in the presence or absence of continuous background noise presented at 35 dB SPL. Target intensity was varied by up to 40 dB. Although the location of the background noise varied (−80° to 80° in increments of 40°), it was always presented from a single location at any given time. Targets and background noise were both broadband with flat spectra (0.5-30kHz).

The authors report the percentage of active neurons for various spatial configurations of target and masker (Furukawa and Middlebrooks, 2001)(Fig. 3), which provides a measure of the population response. We therefore fit Gaussians to these data to generate population tuning curves for different background noise conditions (quiet, or background noise presented at 0°, 40° and 80° away from the midline), and assume symmetric responses for the left hemisphere. We then used these population tuning curves to simulate behavioral data for different background noise conditions. On each simulated trial, we generated left- and right-hemisphere responses for a sound presented at a specific location. This was done by identifying the appropriate neural responses from the population tuning curves, and adding a random term, drawn from a Gaussian distribution, to simulate neural noise. The variance of this Gaussian distribution was chosen so that the overall performance of the model (mean error magnitude) matched that of human participants (either for broadband or narrowband stimuli, although this produced no appreciable difference in the predicted pushing effects). To simulate different background noise conditions, neural responses were generated using the population tuning curves appropriate to each condition.

On each simulated trial, we calculated the difference in response between the two hemispheres. To decode this hemispheric difference, the model then identified the target location that produced the most similar hemispheric difference under quiet conditions. This was then used to generate the behavioral output of the model. When background noise was absent, this led to accurate performance. However, when background noise was present, neural responses were often decoded incorrectly, which led the model to make systematic errors in sound localization.

To better understand the link between changes in population tuning curves and changes in model output, we also ran the model above using population tuning curves that were manipulated in specific ways (i.e. tuning width was sharpened or responses were reduced). In such cases, we started with population tuning curves that were Gaussian fits to real neurophysiological data obtained in the absence of background noise (Furukawa and Middlebrooks, 2001). We then changed the standard deviations of these Gaussians (to sharpen tuning) or multiplied the Gaussians by a scale factor (to reduce responses).

### Code Accessibility

All relevant code is available on request from the corresponding author.

## Results

### Background noise reduces sound localization errors following simulated unilateral hearing loss

We began by measuring the impact of background noise on sound localization under normal hearing conditions, focusing on the role of binaural spatial cues (Interaural Time Differences: ITDs; Interaural Level Differences: ILDs). To do this, we used narrowband target sounds, which prevent subjects from using monaural spectral cues to sound location. Target sounds were presented at various locations in the front hemifield, either in quiet or in the presence of continuous broadband noise located directly in front of the listener (Fig. 1A). This was done to facilitate comparison with previous work (Furukawa and Middlebrooks, 2001). Targets also varied in frequency and intensity (corresponding to different signal-to-noise ratios (SNRs) in the background noise condition; all SNRs ≥ 0), but the effects of background noise were very similar in each case and so are plotted together (no significant interactions between frequency/intensity and noise condition, mixed-effects ANOVA, p > 0.05).

**Figure 1.**
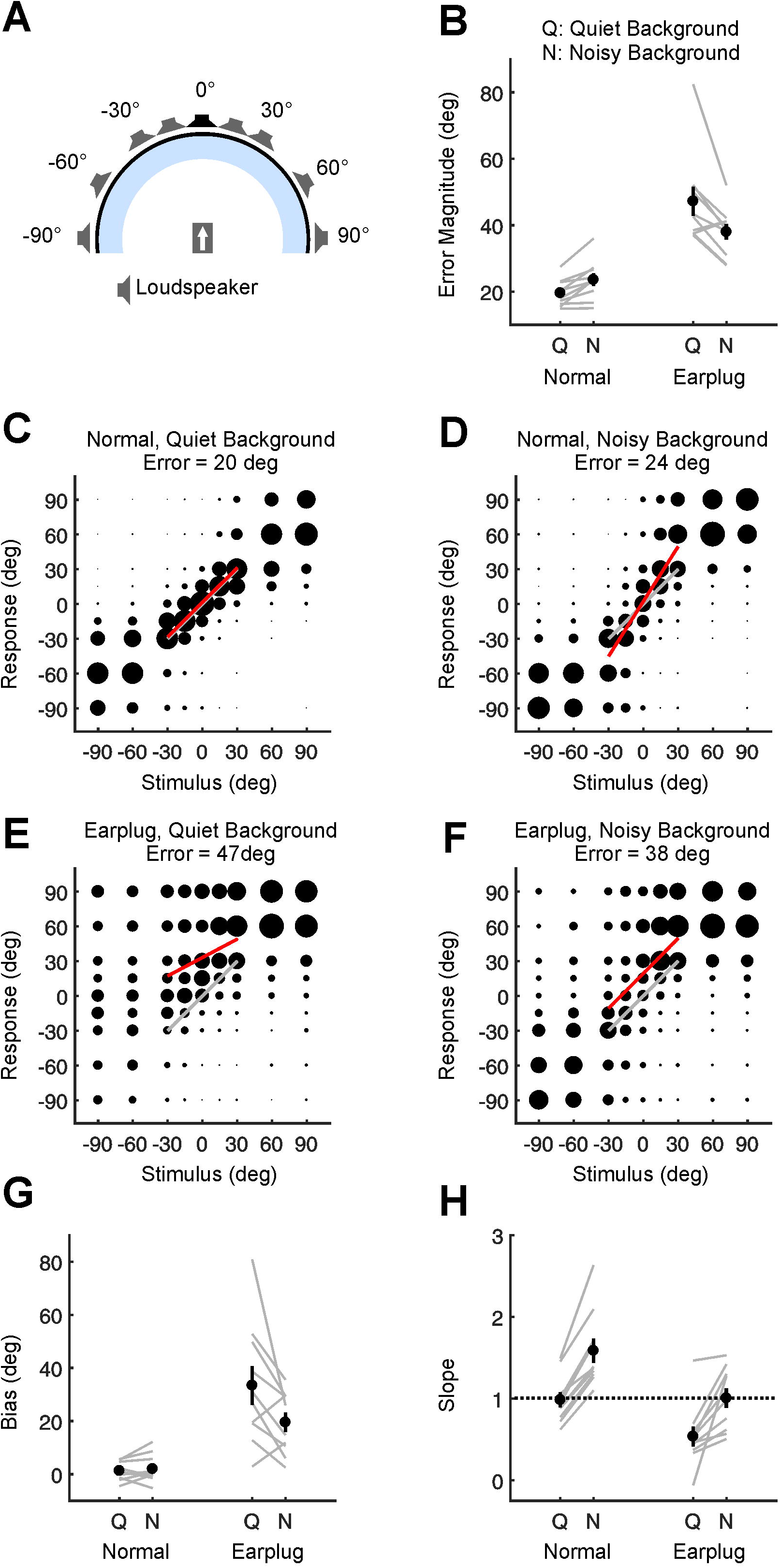
Behavioral effects of background noise on narrowband sound localization. **A** Schematic of speaker array used to test sound localization. On each trial, a target sound was presented from one of the loudspeakers shown. Subjects sat at the centre of the array and used a graphical user interface to indicate their response. To create a noisy background, broadband noise was presented continuously at the midline (0°, black). **B** Magnitude of sound localization errors (black: mean ± SEM; gray: individual subject data) are plotted for different experimental conditions. Subjects located sounds with normal hearing or with an earplug in one ear, and with background noise turned either off (Q: quiet) or on (N: noisy). **C** Localization data for normal hearing condition with background noise off. Joint distribution of stimulus and response is shown averaged across all subjects. Circle size is proportional to the number of trials corresponding to each unique combination of stimulus and response. Results of linear regression (restricted to stimulus locations ±30° away from the midline) are shown for actual data (red) and perfect performance (gray). Mean error magnitude is displayed above main plot. **D-F** Identical to (C) but showing localization data for normal hearing condition with background noise on (D), earplug condition with background noise off (E), or earplug condition with background noise on (F). **G** Identical to (B) but showing subjects’ bias values. Positive values indicate a tendency to mislocalize sounds toward the side of the ear that did not receive an earplug. **H** Identical to (B) but showing slopes of linear regression lines fit to localization data. Perfect performance on task is associated with a slope of 1 (dashed line); values above or below this reflect errors.

With normal hearing, subjects performed this task well in a quiet environment, making relatively small errors (Fig. 1B,C). Performance was worse, however, in the presence of background noise, with subjects making larger errors (p < 0.05, post-hoc test; Fig. 1B,D). We then asked whether individuals are more vulnerable to background noise if they experience a hearing loss in one ear. To do this, we simulated a partial unilateral hearing loss by requiring the same subjects to wear an earplug in one ear. This delays and attenuates the input to the plugged ear (Kumpik et al., 2010; Keating et al., 2016), which alters the two primary sound localization cues (ITDs and ILDs). When subjects wore an earplug, sound localization was considerably impaired relative to normal hearing conditions, irrespective of whether the background noise was off (p<0.05, post-hoc test; Fig. 1B,E) or on (p<0.05, post-hoc test; Fig. 1B,F).

However, when subjects wore an earplug, the effect of background noise was opposite to that observed with normal hearing (significant interaction between background noise and hearing loss conditions; mixed effects ANOVA, F_(1,576)_ = 45.1, P < 0.001; Fig. 1B). Surprisingly, we found that background noise improved sound localization when subjects wore an earplug (p<0.05, post-hoc test; Fig. 1B). This result was not limited to a specific target frequency, with similar results observed if we conducted separate analyses for low-frequency targets (< 1.5 kHz; where ITDs are the primary cue to sound location) or high-frequency targets (> 1.5 kHz; where ILDs are the primary cue to sound location). In each case, background noise improved sound localization following simulated unilateral hearing loss (post-hoc tests, p < 0.05; significant interactions between background noise and hearing loss conditions; mixed effects ANOVAs, Low Frequency Targets: F_(1,144)_ = 5.8, P = 0.018; High Frequency Targets: F_(1,464)_ = 42.5, P < 0.001).

### Background noise improves performance by reducing bias and increasing perceptual discriminability

Previous studies have shown that an earplug impairs sound localization because sounds are mislocalized toward the side of the open ear (i.e. subjects are biased) (Kumpik et al., 2010; Keating et al., 2016). This occurs because the earplug changes ITDs and ILDs in ways that favor the side of the open ear. Localization is also impaired because subjects are less able to discriminate between sounds presented at different locations, which flattens the relationship between stimulus and response (slope of the stimulus-response relationship becomes closer to 0) (Kumpik et al., 2010; Keating et al., 2016). To better understand our results in light of previous work, we therefore used linear regression to estimate bias and slope values for different conditions. To facilitate comparison with previous work (Kumpik et al., 2010), we restricted this analysis to stimulus locations < 45° from the midline (Fig. 1C-F, red lines; see Methods).

Under normal hearing conditions, subjects showed very little bias, with similar bias values observed in the presence and absence of background noise (p > 0.05, post-hoc test, Fig. 1C,D,G). Although an earplug produced a large bias toward the side of the open ear (bias values > 0; p < 0.05, post-hoc tests; Fig. 1E,G), the magnitude of this bias was reduced in the presence of background noise (p < 0.05, post-hoc test; significant interaction between background noise and hearing loss conditions; mixed effects ANOVA, F_(1,576)_ = 18.5, P < 0.001; Fig. 1F,G). In other words, background noise reduced subjects’ tendency to mislocalize sounds toward the side of the open ear following simulated unilateral hearing loss.

If subjects were to perform our task perfectly, the slope of the relationship between stimulus and response would be equal to 1, with deviations above or below this value reflecting errors in sound localization. When locating narrowband sounds in a quiet environment, normal hearing subjects showed slope values very close to 1 (Fig. 1C,H). However, in the presence of background noise, slope values increased (p < 0.05, post-hoc test; Fig. 1D,H). This occurs because the perceived locations of target sounds are pushed away from the location of the background noise (i.e. the midline). For example, when a target sound is presented at 30°, it tends to be perceived further to the right (~50°; Fig. 1D), with symmetric errors observed on the left hand side. This means that background noise increases sound localization errors under normal hearing conditions.

Conversely, when subjects wore an earplug, slope values were < 1 when locating sounds in a quiet environment, and were considerably lower than the slope values observed with normal hearing (p < 0.05, post-hoc test; Fig. 1C,E,H). This is because subjects are less able to distinguish between sounds presented at different locations. However, in the presence of background noise, slope values increased (p < 0.05, post-hoc test; Fig. 1F,H) and became very close to 1. This means that background noise increased slope values similarly for both normal hearing and earplug conditions (no significant interaction between background noise and earplug conditions, mixed effects ANOVA, F_(1,576)_ = 2.8, P = 0.092; Fig. 1H). However, whilst this increase moved slope values away from 1 under normal hearing conditions (post-hoc test, p < 0.05), it moved slope values closer to 1 when subjects were wearing an earplug (post-hoc test, p < 0.05). Consequently, background noise improves narrowband sound localization following simulated unilateral hearing loss by both reducing bias and returning slope values closer to their optimal value (i.e. 1). In this way, background noise partly reverses the two main effects of simulated unilateral hearing loss.

### Broadband sound localization is more robust to the effects of background noise

Since we presented our background noise directly in front of the listener, a simple explanation for a reduction in bias is that the background noise acts as a reference point that informs subjects where the midline is. When wearing an earplug, subjects could therefore use the perceived location of the background noise to estimate how biased they are and shift their localization responses to compensate (note that this could be a useful strategy in everyday life, not only in our task). However, if this were the case, bias would be shifted by the same amount for all target sounds, including those with broadband spectra. To test this, we therefore investigated the effect of background noise on sound localization using broadband targets.

Contrary to this hypothesis, we found that background noise changed bias values (and overall errors) in a stimulus-specific way (i.e. data obtained using broadband targets differed from that obtained using narrowband targets; significant 3-way interactions between background noise, hearing loss and stimulus conditions; mixed-effects ANOVAs; bias values: F_(1,928)_ = 10.9, P = 0.001; overall errors: F_(1,928)_ = 23.6, P < 0.001). In particular, when subjects located broadband sounds, background noise had no effect on bias values or overall errors (p > 0.05 for both normal-hearing and hearing loss conditions, post-hoc tests; Fig. 2A-F). For broadband targets, the effects of background noise were therefore both less detrimental (normal hearing) and less beneficial (plugged hearing) than those observed for narrowband targets. This means that broadband sound localization is relatively robust to the effects of background noise. It also suggests that our subjects are not simply using the background noise as a reference point and therefore requires an alternative explanation.

**Figure 2.**
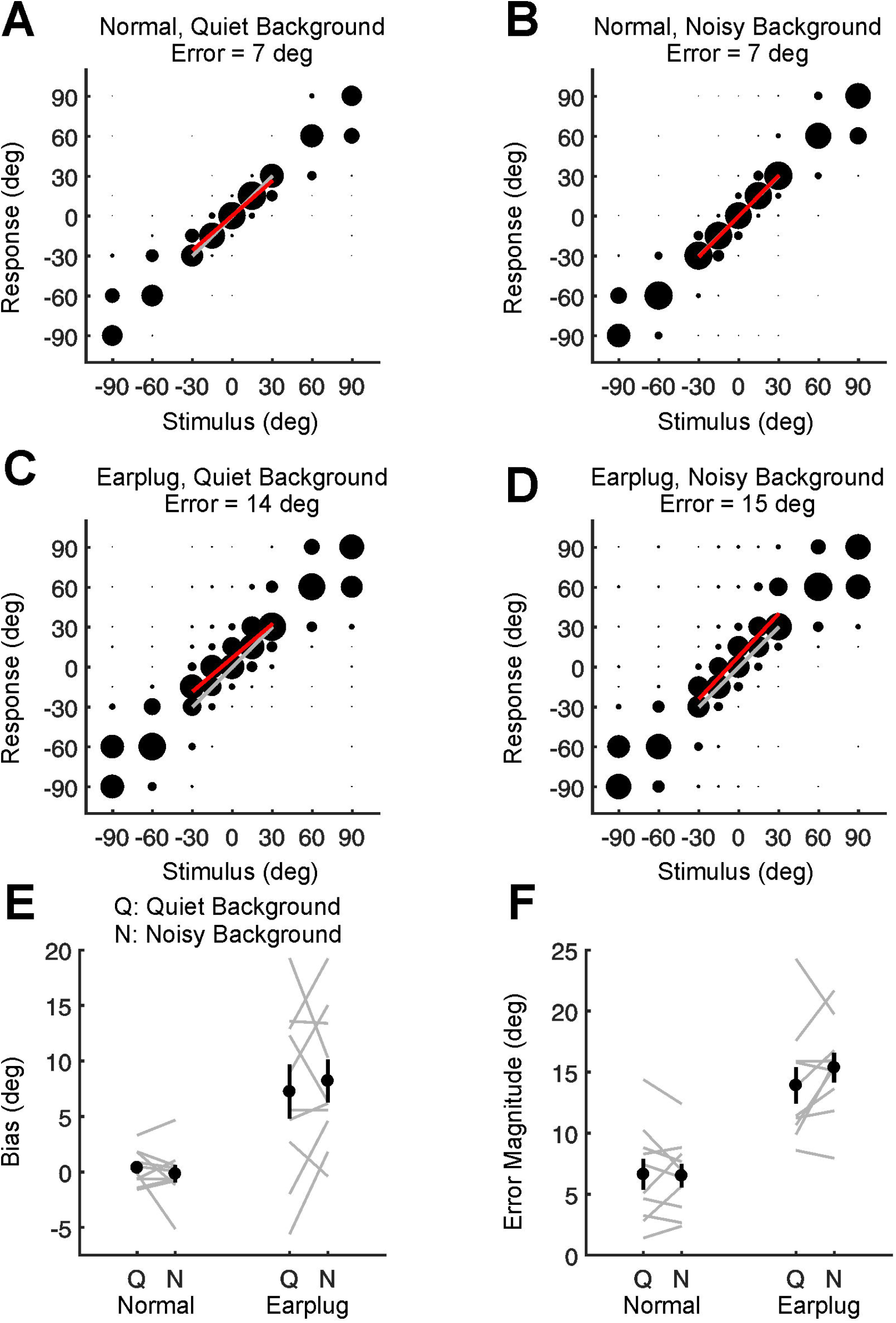
Behavioral effects of background noise on broadband sound localization. **A** Localization data for normal hearing condition with background noise off. Joint distribution of stimulus and response is shown averaged across all subjects. Circle size is proportional to the number of trials corresponding to each unique combination of stimulus and response. Results of linear regression (restricted to stimulus locations ±30° away from the midline) are shown for actual data (red) and perfect performance (gray). Mean error magnitude is displayed above main plot. **B-D** Identical to (A) but showing localization data for normal hearing condition with background noise on (B), earplug condition with background noise off (C), or earplug condition with background noise on (D). **E** Bias values (black: mean ± SEM; gray: individual subject data) are plotted for different experimental conditions. Positive values indicate a tendency to mislocalize sounds toward the side of the ear that did not receive an earplug. Subjects located sounds with normal hearing or with an earplug in one ear, and with background noise turned either off (Q: quiet) or on (N: noisy). **F** Identical to (E) but showing average magnitude of errors made by subjects for different conditions.

### Background noise produces pushing effects in the hemispheric model of sound localization

We next considered previous work that investigated the spatial tuning of auditory neurons (cat A2) for normal hearing conditions whilst continuous broadband noise was presented directly in front of the listener (Furukawa and Middlebrooks, 2001). One key result from this work is that background noise suppresses neuronal responses in a location-specific way. In particular, when sounds are presented in a quiet background, neurons in cat auditory cortex tend to be broadly tuned to sounds presented on the contralateral side of space. This is reflected in the population response for each hemisphere (Fig. 3A; see Methods). However, when continuous background noise is presented at the midline, the population responses to target sounds become more tightly tuned to spatial location (Fig. 3A). Sharper tuning is a robust effect of continuous background noise, and is also observed for single neurons in both A1 (Brugge et al., 1998; Wood et al., 2018) and A2 (Furukawa and Middlebrooks, 2001). In addition, it has been suggested that these effects may partly reflect corresponding changes in the responses of inferior colliculus (IC) neurons (Mokri et al., 2015).

To assess how sharper spatial tuning might influence behavior, we implemented a popular and simple model of sound localization (McAlpine et al., 2001; Grothe et al., 2010; Keating et al., 2015). According to this model (the hemispheric model), sound location is represented by the difference in mean activity between the two hemispheres (hemispheric difference). We therefore calculated hemispheric differences in activity for different background noise conditions (quiet background or continuous background noise presented at the midline; Fig. 3B), using Gaussian fits to previously published neurophysiological data (Fig. 3A; see Methods) (Furukawa and Middlebrooks, 2001).

**Figure 3.**
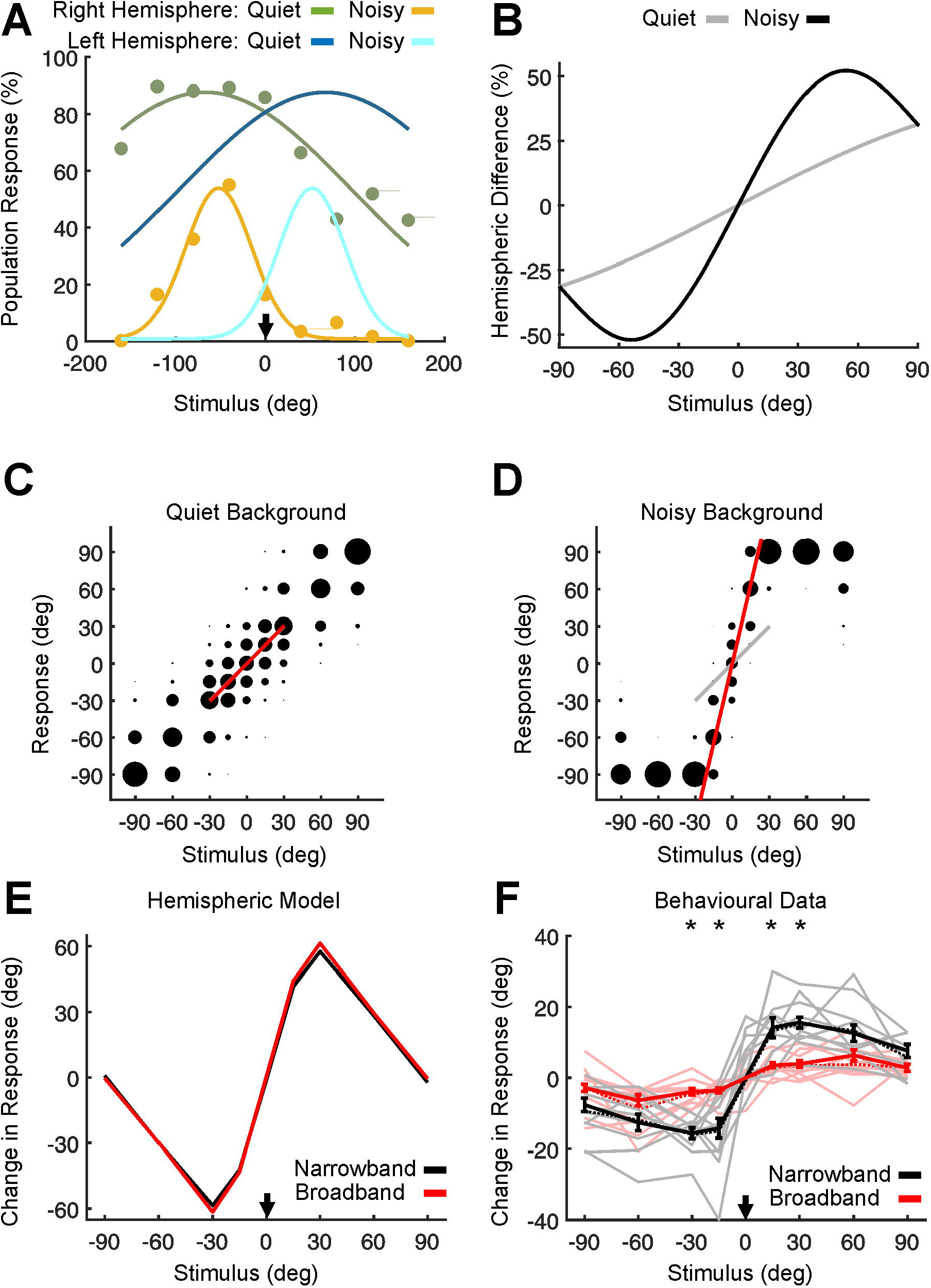
Effect of background noise under normal hearing conditions in the hemispheric model of sound localization. **A** Neural responses to target sounds for left and right auditory cortex either in the presence or absence of continuous background noise located at the midline (0°; arrow). Stimulus locations on the right are represented by positive values. Markers indicate data obtained in cat A2 by Furukawa and Middlebrooks (2001; see also Methods). Lines show Gaussian fits to their data; responses for the left hemisphere are mirrored versions of right-hemisphere responses. **B** Difference in neural response between the left and right hemispheres plotted as a function of target location, calculated using the Gaussian fits in (A). Positive hemispheric differences indicate a greater response in the left hemisphere. Data are shown for conditions in which the background noise was either absent (gray) or present (black). In general, background noise located directly in front exaggerates hemispheric differences in activity, albeit in a location-specific way. **C,D** Sound localization data simulated using the hemispheric differences in (B), either with the background noise off (C) or on (D). For each of these conditions, the joint distribution of stimulus and response is shown. Circle size is proportional to the number of trials corresponding to each unique combination of stimulus and response. Results of linear regression (restricted to stimulus locations ±30° away from the midline) are shown for actual data (red) and perfect performance (gray). Overall errors are constrained to match human behavioral data for narrowband targets. **E** Change in response produced by background noise plotted for each target location. Positive changes in response indicate that background noise shifts responses toward the right; negative changes in response indicate shifts toward the left. Positive target locations are on the right. Data show that target sounds are pushed away from the location of the background noise (midline; arrow). Data are shown for simulations that match overall errors to human behavioral data obtained using narrowband targets (black; computed from data in panels C and D) or broadband targets (red). **F** Identical to (E) except data have been calculated using subjects’ behavioral responses. Data are shown for narrowband targets (black; see also Fig. 1C,D) and broadband targets (red; see also Fig. 2A,B). Asterisks indicate significant differences between narrowband and broadband data (mixed-effects ANOVA, post-hoc tests, p < 0.05). Pale lines show data for individual subjects; dashed lines show data averaged across subjects. Continuous dark lines show data (mean ± SEM) averaged across subjects and hemifields. Close correspondence between continuous and dashed lines indicate that data are symmetric around the midline.

For target sounds presented in the front hemifield background noise exaggerated differences in activity between left and right auditory cortex, particularly for locations ~45° from the midline (Fig. 3B). For example, when a target sound was presented 15 degrees to the right in the presence of background noise, it produced the same population response (i.e. difference in activity between two hemispheres) as a target sound presented further to the right in the absence of background noise. Symmetric effects were observed for sounds located on the left. In the model, background noise is therefore expected to push target sounds away from the midline (i.e. the location of the background noise) relative to quiet conditions.

To illustrate this, we simulated neural responses for quiet and noisy conditions using the hemispheric responses in Fig. 3A. Trial-to-trial variability was simulated by adding a random term to the neural responses on each trial (constrained to match the overall errors observed for our behavioral data). These neural responses were then decoded by the model (see Methods). When we simulated quiet conditions, the model performed well (Fig. 3C). However, when we simulated noisy conditions, the ‘perceived’ (i.e. decoded) locations of target sounds were pushed away from the midline (Fig. 3D). This increased the slope of the relationship between stimulus and response (Fig. 3D, red line), but had no effect on overall bias. This is broadly consistent with what we observed in our behavioral data for narrowband sounds under normal hearing conditions.

### Pushing effects depend on spatial separation between target and background noise

In the model, we next tested the effect of background noise for each target location. To do this, we calculated the mean response for each target location and assessed how much it changed in the presence of background noise (Fig. 3E). In the model, pushing effects were symmetric around the midline (i.e. in each hemifield, target sounds were pushed away from the midline). The greatest pushing effect also occurred for intermediate separations between target and background noise (±30 deg), and declined for greater separations. To test these predictions, we therefore reanalysed our normal-hearing behavioral data to estimate the degree of pushing observed for targets presented at different locations.

In our behavioral data, we found that the pushing effect is also symmetric around the midline (no significant main effect or interaction effects of hemifield; mixed-effects ANOVA, p > 0.05; Fig. 3F). We additionally found that the pushing effect is greatest for intermediate separations between target and background noise (30 deg for narrowband sounds, 60 deg for broadband sounds), and declines for greater separations. However, narrowband targets showed greater pushing effects than broadband targets, particularly close to the midline (significant differences between narrowband and broadband data at 15° and 30°; post-hoc tests, p < 0.05). The pushing effect for narrowband targets was also more sensitive to the separation between target location and background noise (relative to broadband targets; significant interaction between stimulus type and target eccentricity; mixed-effects ANOVA, F_(3,152)_ = 2.7, P = 0.048).

### Sharper spatial tuning produces pushing effects that vary with target location

In the neurophysiological data, background noise reduces the magnitude of neural responses and sharpens their spatial tuning (Fig. 3A) (Furukawa and Middlebrooks, 2001). To isolate the role of each of these factors, we therefore used the hemispheric model to simulate what happens when we vary each of these factors separately. When we reduced the magnitude of the neural responses, but kept the tuning widths constant (Fig. 4A), the hemispheric differences in activity were reduced. In the hemispheric model, the ‘perceived’ locations of target sounds were therefore pulled toward the midline (Fig. 4B), which is opposite to what we observed in our behavioral data (Fig. 3F).

**Figure 4.**
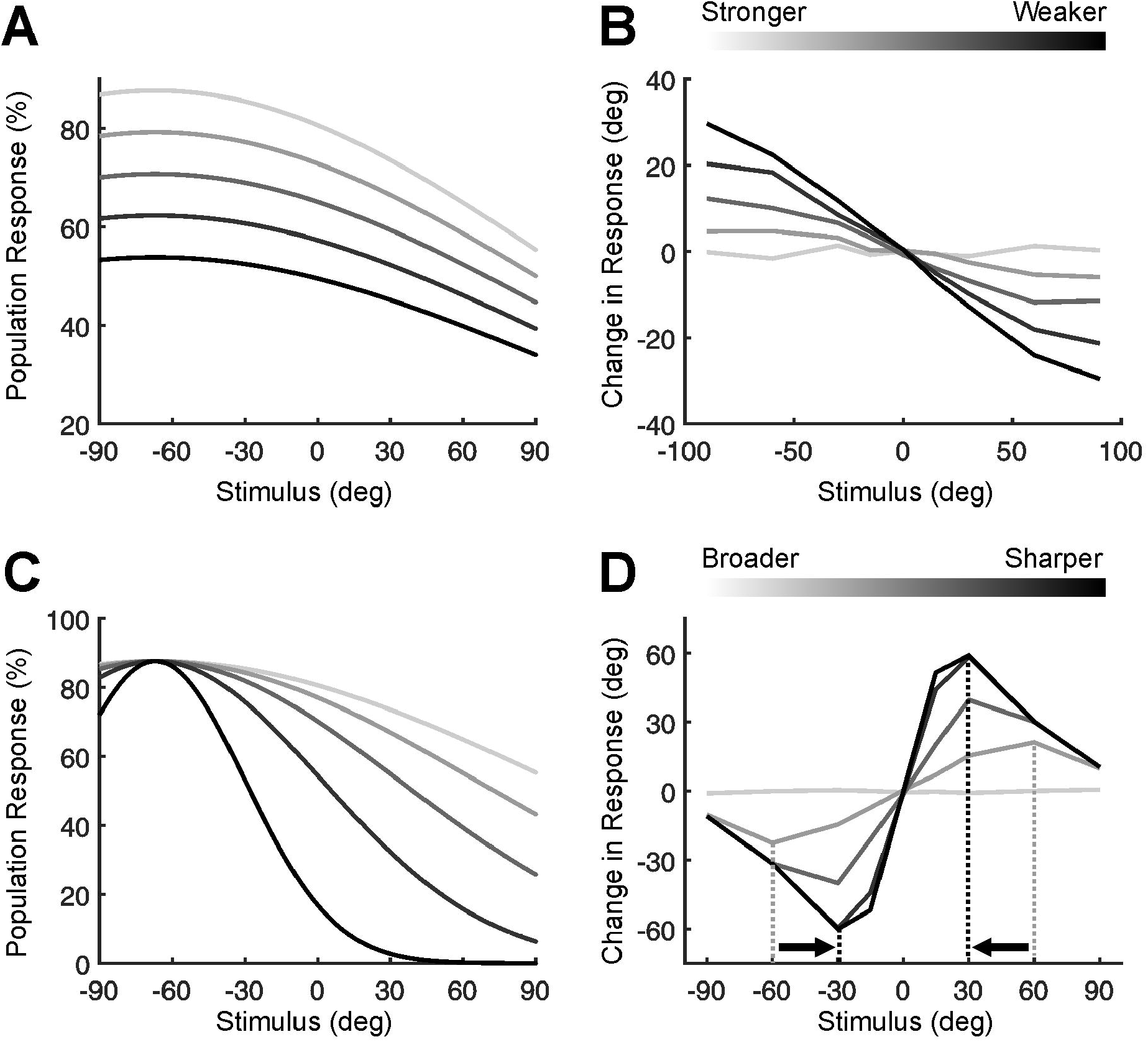
Dissociable effects of sharper tuning and reduced responsiveness in the hemispheric model. **A** Population responses for left hemisphere plotted as a function of stimulus location. Positive stimulus values correspond to locations on the right. Lightest gray shows Gaussian fit to real neural responses obtained in quiet (identical to Fig. 3A). These responses can be reduced artificially using a scaling factor that keeps the tuning width constant. Progressively greater reductions in neural response correspond to progressively darker shades. For clarity, data are shown for right hemisphere only; left-hemisphere data are symmetric around the midline. **B** Reducing neural responses produces changes in the behavioral output of the hemispheric model. Changes in model output (relative to baseline) are plotted as a function of stimulus location. Positive target locations are on the right. Positive changes indicate that the model ‘perceives’ (i.e. decodes) target sounds further to the right (relative to baseline). Overall, reducing neural responses pulls the ‘perceived’ location of target sounds toward the midline, particularly for peripheral sounds. **C** Identical to (A) except the tuning of neural responses has been progressively sharpened whilst keeping the maximum neural response constant. Darker shades correspond to sharper tuning curves. **D** Identical to (B) but showing the impact of sharper tuning on model output. Overall, sharper neural tuning (darker shades) pushes the ‘perceived’ location of sounds away from the midline, particularly for target sounds that are located intermediate distances from the midline (for comparison, see also Fig. 3 E,F). As tuning width is progressively sharpened, the target location associated with the greatest change in response shifts toward the midline (arrows).

However, when we sharpened the spatial tuning of the neural responses, but kept their maxima constant (Fig. 4C), the hemispheric model produced pushing effects (Fig. 4D) that are broadly similar to those observed behaviorally (Fig. 3F). As the tuning width becomes sharper, the model predicts an increase in the magnitude of the pushing effect, particularly for targets located close to the midline. Consequently, the model predicts a relationship between the magnitude of the pushing effect and the target location that exhibits the greatest amount of pushing. Interestingly, this relationship parallels the differences we observed in our behavioral data between narrowband and broadband targets (Fig. 3F). This means that the stimulus differences we observed in our behavioral data are consistent with differences in a single parameter (i.e. degree to which spatial tuning is sharpened by background noise).

### Effects of lateralized background noise in the hemispheric model

Previous neurophysiological studies have not investigated the effects of background noise following unilateral hearing loss. However, researchers have shown changes in the spatial tuning of neurons if they are exposed to background noise presented away from the midline (Fig. 5A) (Furukawa and Middlebrooks, 2001). In such circumstances, the background noise produces binaural spatial cues (ITDs and ILDs) that favor one side of space. Under these conditions, the balance of neural activity between the two hemispheres in response to target sounds is altered. For example, when background noise is presented 40° to the right of the midline, a target located in front of the listener produces a hemispheric difference equivalent to that produced by a target on the left in a quiet environment (Fig. 5B). In the model, the ‘perceived’ location of target sounds is therefore pushed away from the side of the background noise. Similar changes in the hemispheric difference are observed if background noise is presented further from the midline (80°; Fig. 5B,C).

**Figure 5.**
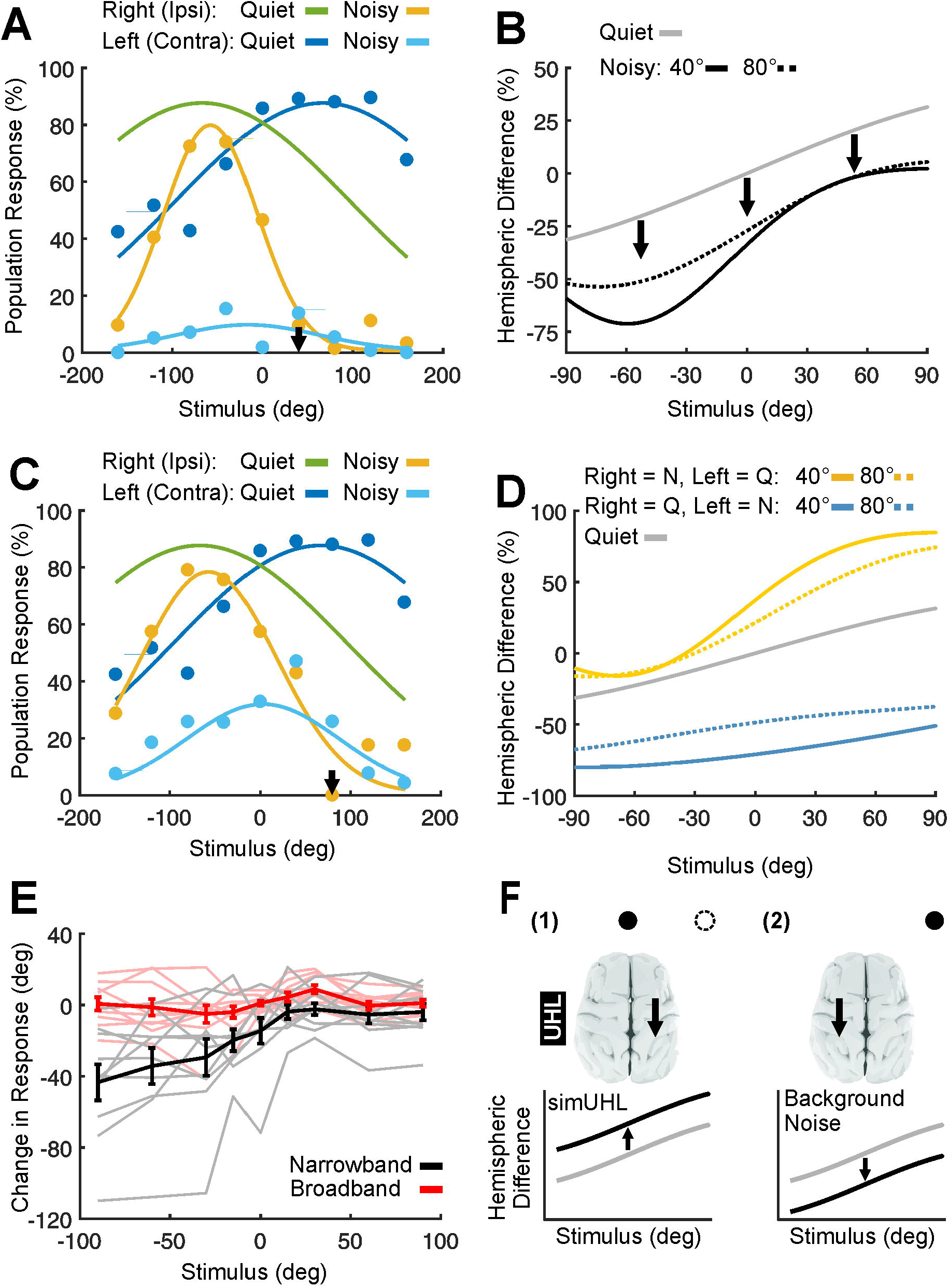
Effects of lateralized background noise. **A** Neural responses to target sounds in the presence or absence of continuous background noise to the right of the midline (40°; arrow). Data are shown for the left hemisphere (contralateral to background noise) and right hemisphere (ipsilateral to background noise). Stimulus locations on the right are represented by positive values. Markers indicate data obtained from cat A2 by Furukawa and Middlebrooks (2001). Lines show Gaussian fits to their data; responses for the left hemisphere are mirrored versions of right-hemisphere responses. **B** Difference in neural response between the left and right hemispheres plotted as a function of target location, calculated using the Gaussian fits in (A). Positive hemispheric differences indicate a greater response in the left hemisphere. Data are shown for conditions in which the background noise was absent (gray), presented 40° to the right (continuous black), or 80° to the right (dashed). **C** Identical to (A) but showing data for background noise located 80° to the right of the midline. **D** Identical to (B), but shows what happens if we simulate the impact of background noise for the right hemisphere only (yellow; hemisphere ipsilateral to the background noise) or left hemisphere only (blue; contralateral to the background noise). Data are shown for background noise located 40° (continuous coloured lines) or 80° (dashed lines) to the right of the midline, as well as for the quiet condition (gray). Effects of background noise in the two hemispheres oppose one another (shift in opposite directions relative to gray line) and partly cancel each other out when the hemispheric difference is computed. **E** Behavioral effects of background noise following simulated unilateral hearing loss for different target locations. Change in response produced by background noise is plotted for each stimulus location. Negative stimulus locations are on the side of the plugged ear. Negative changes in response indicate that background noise shifts responses toward the side of the plugged ear; positive changes in response indicate shifts toward the side of the unoccluded ear. Data are shown for narrowband (black) and broadband (red) targets. Pale lines show data for individual subjects. Continuous dark lines show data (mean ± SEM) averaged across subjects. **F** Hypothesis to explain beneficial effects of background noise following simulated unilateral hearing loss. (1) when a simulated unilateral hearing loss (simUHL) is induced in the left ear, responses to target sounds are reduced in the right hemisphere (middle; Keating et al., 2015), which alters the balance of activity between the two hemispheres (bottom). Background noise presented at the midline (filled black circle, top) is also perceived on the right (dashed circle) because it produces binaural cues that favor that side. (2) When background noise is presented on the right (filled black circle, top), responses to target sounds are reduced in the left hemisphere (middle; Furukawa and Middlebrooks, 2001), which alters the balance of activity between the two hemispheres (bottom; opposite direction to simUHL). Background noise may therefore help rebalance activity between the two hemispheres following simUHL.

Interestingly, when background noise is presented on one side of space, the population response in each hemisphere is affected in a different way. For example, in the hemisphere ipsilateral to the background noise, spatial tuning is primarily sharpened by background noise whilst the peak response remains relatively unchanged (Fig. 5A,C). Conversely, in the hemisphere contralateral to the background noise, the peak response is considerably reduced by background noise whilst the tuning curve width remains less affected (Fig. 5A,C). To illustrate the role played by each hemisphere in the hemispheric model, we therefore considered what would happen to the hemispheric difference if background noise affected only a single hemisphere (i.e. we calculated the hemispheric difference using spatial tuning curves obtained in the presence of noise for one hemisphere, and in quiet for the other hemisphere). If we simulate the effects of background noise solely for the hemisphere contralateral to the background noise, the hemispheric difference shifts toward more negative values (blue, Fig. 5D). This is consistent with target sounds being pushed away from the side of the background noise. By contrast, if we simulate the effects of background noise solely for the hemisphere ipsilateral to the background noise, the hemispheric difference shifts toward more positive values (yellow, Fig. 5D). This is consistent with target sounds being pulled toward the side of the background noise. When the hemispheric difference is computed, the effects of lateralized background noise in each hemisphere therefore oppose one another and partly cancel.

### Background noise produces greater pushing effects on the side of the earplug

If subjects wear an earplug, background noise presented at the midline produces binaural spatial cues that favour the side of the open ear(Eric Lupo et al., 2011; Keating et al., 2016). Consequently, subjects perceive the background noise on the side of the open ear (all subjects reported this, but it is also evident in their responses to identical stimuli of short duration; Fig. 2 E). In light of the modelling results above, we might expect background noise to push target sounds away from the side of the open ear, which is the side on which the background noise is perceived. For narrowband sounds, this is precisely what we found (Fig. 1G). However, we wanted to know whether this pushing effect varies with target location. We therefore reanalysed our behavioral data for subjects wearing an earplug.

In particular, we calculated the mean behavioral response for each target location and assessed how much it changed in the presence of background noise (Fig. 5E). Although background noise pushed narrowband targets away from the side of the open ear (change in response < 0), this pushing effect was greater on the side opposite the open ear (significant effect of hemifield, post-hoc test, p < 0.05). However, very different effects of background noise were observed for broadband targets (significant interaction between stimulus type and hemifield, mixed-effects ANOVA, F_(1,144)_ = 41.8, p < 0.001), particularly on the side opposite the open ear (significant effect of stimulus type on that side; post-hoc test, p < 0.05; Fig. 5E). This is primarily because background noise had very little effect on broadband localization.

## Discussion

Many auditory abilities are impaired in the presence of background noise, particularly in individuals who suffer from hearing loss (Moore, 1996; Lorenzi et al., 1999a; Bronkhorst, 2000; Lorenzi et al., 2006; Helfer and Freyman, 2008; Akeroyd, 2014). Surprisingly, however, we found that background noise improved narrowband sound localization when subjects experienced a simulated hearing loss in one ear. By contrast, localization of broadband sounds was more robust, and was less affected by either simulated hearing loss or background noise. A simple neurophysiological model also provided insight into how the behavioral effects of background noise might arise.

### Background Noise Distorts Sound Localization Under Normal Hearing Conditions

Under normal hearing conditions, sound localization was relatively unaffected by background noise, with errors only increasing for narrowband targets. However, we ensured that target sounds were clearly audible by using signal-to-noise ratios (SNRs) ≥ 0. Previous work suggests that background sounds impair sound localization by reducing target audibility (Good and Gilkey, 1996; Abouchacra et al., 1998; Lorenzi et al., 1999a, b; Brungart et al., 2005; Kopco et al., 2010; Kerber and Seeber, 2012; Lingner et al., 2012; Wood and Bizley, 2015). Greater effects of background noise may therefore occur at more adverse SNRs. Nevertheless, we found that the perceived location of targets was pushed away from the location of the background noise. Although previous studies have observed ‘pulling’ effects between sounds that are grouped together (Lee et al., 2009), our target sounds were clearly distinct from the background noise, which tends to produce ‘pushing’ effects (Suzuki et al., 1993; Canevet and Meunier, 1996; Getzmann, 2002; Best et al., 2005; Reed and van de Par, 2015). We also found greater pushing effects for narrowband targets, which indicates that the pushing effect is a feature of binaural spatial processing. Previous studies have shown that the pushing effect increases with spatial separation between target and background noise (Best et al., 2005). However, because we tested a wider range of spatial separations, we found that the pushing effect declines for even greater spatial separations. By using broadband and narrowband targets, we also found that the effect of spatial separation is stimulus-specific.

To explain the behavioral effects of background noise, previous models have relied upon a neural map of space with an array of neurons (or ‘spatial channels’) tuned to different locations (Suzuki et al., 1993; Best et al., 2005). In these models, sound location is represented by the neurons (channels) that fire most, and competing sounds repel one another because of competitive interactions between neurons tuned to adjacent locations in space. These models therefore predict greater pushing effects for targets that are closer to the background noise, which is not observed behaviorally (Best et al., 2005). A second difficulty for these models is that neurophysiological studies in mammals are inconsistent with an array of neurons tuned to different spatial locations. Instead, neurons within a single hemisphere tend to be broadly tuned to sounds presented in the contralateral hemifield(Brugge et al., 1998; Furukawa and Middlebrooks, 2001; McAlpine et al., 2001; Grothe et al., 2010; Mokri et al., 2015). This has led to the suggestion that sound location is represented by the difference in activity between the two hemispheres (hemispheric model) (McAlpine et al., 2001; Grothe et al., 2010). When applied to data recorded from cat A2 (Furukawa and Middlebrooks, 2001), the hemispheric model captured key features of our behavioral data. This includes a predicted relationship between the magnitude of the pushing effect and the target location that exhibits the greatest amount of pushing. In the model, these effects occur because background noise sharpens spatial tuning, which has been observed in a number of different studies (Brugge et al., 1998; Furukawa and Middlebrooks, 2001; Mokri et al., 2015; Wood et al., 2018).

Nevertheless, the greater robustness of broadband sound localization (relative to narrowband) suggests that the brain may do more than simply average the activity of neurons tuned to different frequencies (Day and Delgutte, 2013; Goodman et al., 2013). If this is the case, then the effects of background noise may be smaller in neurons that integrate information across frequency. One implication of this is that spatial representations may become more robust at higher levels of the auditory system, where frequency tuning tends to be broader. Greater robustness may also be achieved by taking into account differences in the spatial selectivity of individual neurons (Stecker et al., 2005; Keating et al., 2015), or by relying on a sub-population of noise-robust neurons (Mokri et al., 2015). However, our results suggest that the effects of background noise are not entirely eliminated, even when target sounds are clearly audible.

### Beneficial Effects of Background Noise Following Simulated Unilateral Hearing Loss

Consistent with previous studies, we found that sound localization was impaired when subjects experienced a hearing loss in one ear (Slattery and Middlebrooks, 1994; Wightman and Kistler, 1997; Hawley et al., 1999; Van Wanrooij and Van Opstal, 2004, 2007; Kumpik et al., 2010; Irving and Moore, 2011; Strelnikov et al., 2011; Rothpletz et al., 2012; Agterberg et al., 2014; Keating et al., 2016; Parisa et al., 2017; Asp et al., 2018; Nelson et al., 2018). This disruption of sound localization was greater for narrowband sounds (relative to broadband), which suggests that the effects of unilateral hearing loss may be mitigated by combining information across frequency. Previous work has shown that subjects rely more on the spectral cues provided to the intact ear following unilateral hearing loss (Van Wanrooij and Van Opstal, 2004, 2007; Kumpik et al., 2010; Keating et al., 2013; Keating et al., 2016). However, these spectral cues are unavailable if narrowband sounds are used, which would explain worse localization performance for these stimuli. Nevertheless, when subjects wore an earplug, they were still able to locate narrowband sounds, albeit less well. Since this requires subjects to compare the input to the two ears, it indicates that subjects used the residual input to the occluded ear. We would therefore expect greater disruption of sound localization following more severe forms of unilateral hearing loss (Wightman and Kistler, 1997; Van Wanrooij and Van Opstal, 2004; Agterberg et al., 2014; Firszt et al., 2015; Asp et al., 2018).

Although many auditory abilities are impaired by background noise following hearing loss (Moore, 1996; Lorenzi et al., 1999a; Bronkhorst, 2000; Lorenzi et al., 2006; Helfer and Freyman, 2008; Akeroyd, 2014), we found that background noise improved narrowband sound localization following simulated unilateral hearing loss. Since similar results were observed for both low- and high-frequency targets, which respectively rely on ITDs and ILDs (Middlebrooks and Green, 1991), this beneficial effect appears to be a general feature of binaural processing. However, background noise had much less effect on broadband targets. This is consistent with our normal-hearing data, which means that broadband sound localization is more immune to both the negative (normal hearing) and positive (plugged hearing) effects of background noise. Although we investigated what happens when targets and background noise overlap in time, auditory spatial processing can adapt to sounds that precede a target sound but do not overlap (Dahmen et al., 2010; Maier et al., 2012; Stange et al., 2013; Phillips, 2014; Kopco et al., 2017; Tolnai et al., 2017; Ferger et al., 2018). However, previous work has shown that adaptation cannot fully explain the effects of concurrent background noise on sound localization (Best et al., 2005), which points toward additional mechanisms. Similarly, whilst background noise can improve the audibility of subthreshold sounds via stochastic resonance (Zeng et al., 2000; Itzcovich et al., 2017), we saw similar results across a wide range of SNRs, which argues against this.

To understand the beneficial effects of background noise at a neurophysiological level, new studies will be necessary. However, previous work has shown that simulated unilateral hearing loss changes the balance of target responses between the two hemispheres (Keating et al., 2015). When background noise (or preceding sounds) are presented on one side of space, they also alter the balance of target responses between the two hemispheres (Furukawa and Middlebrooks, 2001; Dahmen et al., 2010). Additionally, when subjects wore an earplug, the background noise produced binaural cues that favored the side of the open ear (and was perceived on that side). Consequently, a key hypothesis for future work is that background noise helps rebalance activity between the two hemispheres following unilateral hearing loss (Fig. 5F). When subjects wear an earplug for prolonged periods of everyday life, they show no adaptation to the abnormal binaural cues when tested in quiet environments (Kumpik et al., 2010). This is surprising because subjects can learn to do this if given appropriate training (Keating et al., 2016). However, our results show that subjects rapidly reinterpret abnormal binaural cues in the presence of background noise. If subjects rely on this to maintain accurate sound localization following earplugging, they might not show binaural adaptation in quiet environments. Consequently, whilst the enormous diversity of environmental sounds may affect perception and adaptation in different ways, our results suggest that background noise may be part of the solution as well as the problem.

## Acknowledgements

PK was supported by a Rosetrees UCL Excellence Fellowship, a Wellcome Trust / Academy of Medical Sciences Springboard Award [SBF002\1115] and the NIHR UCLH BRC Deafness and Hearing Problems Theme.

